# Identifying Informative Gene Modules Across Modalities of Single Cell Genomics

**DOI:** 10.1101/2020.02.06.937805

**Authors:** David DeTomaso, Nir Yosef

## Abstract

Two fundamental aims that emerge when analyzing single-cell RNA-seq data are that of identifying which genes vary in an informative manner and determining how these genes organize into modules. Here we propose a general approach to these problems that operates directly on a given metric of cell-cell similarity, allowing for its integration with any method (linear or non linear) for identifying the primary axes of transcriptional variation between cells. Additionally, we show that when using multimodal data, our procedure can be used to identify genes whose expression reflects alternative notions of similarity between cells, such as physical proximity in a tissue or clonal relatedness in a cell lineage tree. In this manner, we demonstrate that while our method, called *Hotspot*, is capable of identifying genes that reflect nuanced transcriptional variability between T helper cells, it can also identify spatially-dependent patterns of gene expression in the cerebellum as well as developmentally-heritable expression signatures during embryogenesis.

## Introduction

Transcriptome-scale profiling at a single-cell resolution has enabled comprehensive categorization of cell types and states in diverse tissues [1, 2, 3], investigation of developmental transitions [4, 5], and in-depth characterization of disease processes [6, 7]. While initial studies focused on the estimation of the transcriptomes in small numbers of cells, the field is rapidly increasing the number of cells per assay [8].

A primary problem that emerges in the analysis of single-cell RNA-seq (scRNA-seq) data is the identification of genes that vary in an informative manner and the organization of these genes into gene modules. A general way to approach this problem is to select for the genes which best reflect the primary axes of variation between cells. Or alternately, finding genes that have similar expression in cells that are close to each other in terms of some biologically-relevant metric. While the most immediate way of defining such a metric is through the use of overall transcriptional similarity between cells, the emergence of multimodal single-cell technologies (which in addition to the cell transcriptome, can profile cellular DNA [9], cell surface proteomes [10], spatial position [11, 12], epigenetic modifications [13], or cell lineages [14]) allows for metric definitions which use other cellular properties. For example, a spatial-metric could be used to compare cells based on physical proximity, or a lineage-metric to compare cells based on clonal relationships. The use of diverse metrics naturally leads to new and interesting questions such as “which genes’ transcription may be influenced by a cell’s location in a tissue?” or “which genes are expressed in a manner consistent with the developmental and clonal relatedness between the cells?”

To facilitate this type of analysis, we have developed a method for gene selection and module identification, called *Hotspot*, which is flexible with respect to the choice of cell metric. Here we present this method and demonstrate its use on three different types of cell metrics - transcriptional, spatial, and lineage - using publicly available and simulated single-cell data.

With transcriptional-only data, we show that our procedure selects informative genes and gene components in a sample of CD4+ T cells, and we demonstrate favorable performance when compared with other methods of feature selection. Notably, our procedure operates on the output of single-cell dimensionality reduction procedures, and is particularly useful when combined with emerging non-linear procedures (such as SIMLR [15], scVI [16], or DCA [17]) for which a direct association between genes and model components is not easily computed. Using a spatial-metric, we demonstrate *Hotspot’s* use for identifying spatial patterns of transcription in mouse cerebellum [12] and benchmark against a leading method, SpatialDE [18], showing similar performance with significantly reduced computation time. We further use a lineage-metric to show how our procedure is able to identify developmentally-associated gene modules using a CARISP/Cas9-based lineage-tracing data during mouse embryogenesis [14]. To our knowledge, *Hotspot* is the first method proposed for this purpose.

*Hotspot*, is implemented as an open-source Python package and is available for use at http://www.github.com/Yoseflab/Hotspot.

## Results

### Leveraging Similarity Maps for Feature Selection and Module Identification

The Hotspot procedure is divided into two main steps. In the first step, a feature selection procedure is performed to isolate the genes which exhibit nonrandom patterns of expression within the similarity map. This is followed by the second step in which correlations are evaluated between genes (using the similarity map) and genes are grouped into modules.

To compute the similarity graph, some notion of ‘similarity’ between cells is needed. If identifying spatial patterns, similar cells would be nearby cells in 2 or 3-dimensional space. For lineage data, similar cells would be nearby cells within an inferred lineage tree [14]. The similarity map can also come from the expression data itself for the case of identifying transcriptional modules in a cluster-free manner. Here similar cells are cells with similar overall transcriptional profiles - i.e. nearby cells in the reduced dimensional space that is output from modeling procedures such as principal components analysis (PCA), diffusion maps [19], or other factor models such as ZINB-WaVE [20], scVI [16], or DCA [17].

Once a notion of similarity is selected, a similarity graph can be computed between cells as a K-nearest neighbors graph. In this graph representation, each cell is a node, and edges connect each cell to the K (configurable, but typically around 30) most similar cells. This structure is already frequently used in single-cell analysis for clustering [21] and visualization [22].

For the feature selection step, we seek genes whose expression is well represented by the similarity graph - genes for which a cell’s expression is highly predictable by it’s local neighborhood. This includes genes which are highly expressed in a particular region of the graph (regardless of whether this region falls naturally into a cluster) or genes whose expression exhibits a smooth gradient across the space. To quantify this, we make use of a test statistic which extends previous work in demographic analysis [23] and machine learning [24] by incorporating a parametric null model for each gene’s expression so that statistical significance and effect size can be efficiently estimated. These metrics are then used to rank and filter genes both to aid in exploratory analysis and to limit the scope of the module identification problem.

In the second step, the genes selected in the first step are grouped into modules based on co-expression. Here, we extend the test statistic used in the feature selection step to operate on pairs of genes and quantify the degree to which two genes have correlated expression in the same regions of the similarity map. This quantifies, for example, whether cells expressing high levels of one gene tend to be near cells expressing high levels of another gene. After this metric is evaluated on all pairs of genes, a hierarchical clustering procedure is used to group genes into modules. For full details on this procedure as well as the feature selection method, see the Methods section.

### Uncovering transcriptional modules in CD4+ T Cells

As a first evaluation for this approach, we simulated single-cell RNA-seq libraries with SymSim [25] so that we could compare *Hotspot’s* prediction against a known, ground truth. For this test, five distinct transcriptional modules were simulated in 3000 cells with varying effect sizes per gene along with a set of negative control genes that varied independently. To mimic the complexity of real data, we simulated a module structure which includes both nested structure and intersecting components(Supplementary Figure 1A). We first evaluated the feature-selection procedure to determine how well Hotspot could detect the 500 genes which participated in the simulated modules and distinguish these from the remaining 4500 genes which vary independently. Compared with the highly-variable genes procedure (”HVG”, as implemented in Seurat [26]), and the NBDisp procedure from [27], Hotspot is able to attain significantly improved performance as demonstrated by Precision-Recall curves of Figure 1B. Additionally, Hotspot shows marginal improvment over a PCA-based feature selection described in [27] (and in the Methods section).

**Figure 1:**
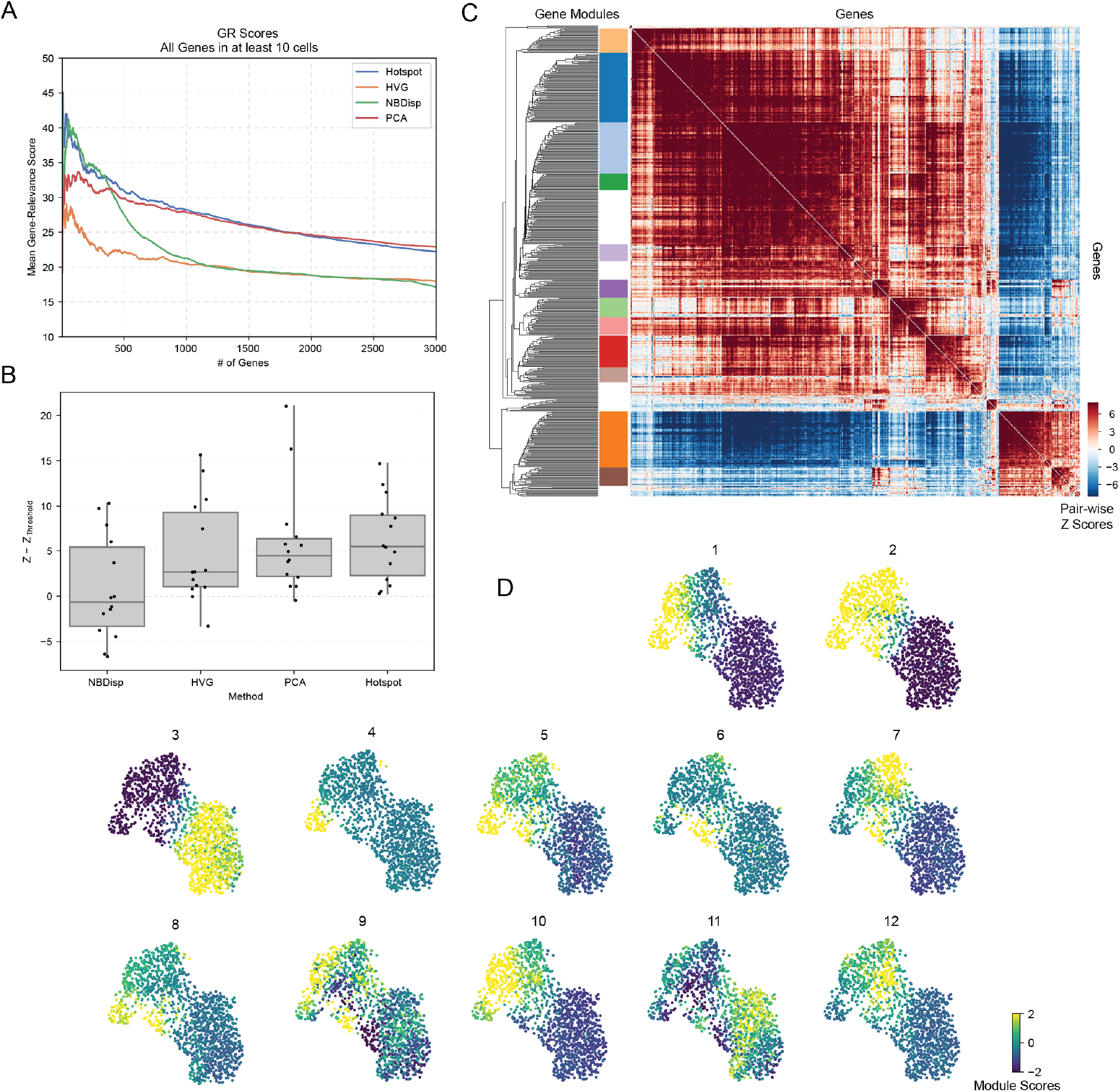
Evaluating Hotspot on a Transcriptional Dataset of CD4 T Cells. A) Comparing feature selection for four methods - Hotspot, HVG (highly variable genes, as implemented in Seurat[26]), NBDisp (from [27]), and a PCA-based procedure (as described in [27]). For each method, the average Gene Relevance score (computed as the number of CD4-relevant gene sets from MSigDB [28] a gene is found in) is computed on the output set of genes as the selection threshold is varied. B) For each method in (A) the top 1000 genes were used to build a latent space model in scVI [16]. The autocorrelation Z-scores of 15 surface protein expression (not included in model building) are used to evaluate estimates of cell state. Shown is the difference (change in Z-score) for each of the 15 proteins when comparing the scVI models from the selected top 1000 features to a model built from all genes above an expression threshold (1̃2000). C) Top 500 genes selected by Hotspot are grouped into 12 modules on the basis of pair-wise local correlation. D) Module summary scores are visualized on a UMAP of the dataset.

To evaluate the module assignment procedure in *Hotspot*, independently of feature selection, we pre-selected the 500 simulated module genes and then compared gene-module assignment accuracy using either standard correlation or Hotspot’s local graph correlation metric followed by hierarchical clustering. Here we demonstrate that *Hotspot* is able to leverage the similarity graph to more accurately make gene-module assignments than standard measures of correlation (Figure 1C). While improvements are small for genes with the largest simulated gene-effect coefficients, the difference is more pronounced for those with a weaker effect size suggesting that Hotspot is able to effectively leverage the neighborhood graph to help estimate these more difficult cases.

To further demonstrate *Hotspot*, we ran our procedure on a set of 1500 CD4+ T cells filtered from a sample of human PBMCs made available by 10x Genomics. To evaluate feature selection on these cells, we utilized two approaches. We first evaluated our methods’s ability to prioritize biologically relevant genes. To quantify biological relevance, we computed a “gene relevance” (GR) score for every gene, as the number of CD4+ T Cell-related gene sets from MSigDB [28] within which the gene is found (Methods). With this metric, we evaluated the relevance of a set of genes as the average GR score for genes in the set. Under this evaluation, *Hotspot* outperforms the commonly used highly-variable genes selection procedure across all thresholds (Figure 1) and performs better than a PCA-based procedure (Methods) for the top 1800 genes. When comparing against the NBDisp [27] procedure, performance is equivalent for the top few hundred genes after which Hotspot’s genes have consistently higher GR scores.

To evaluate the feature selection procedure in an unsupervised manner, we attempted to quantify the quality of the similarity map generated by scVI. Here we reason that a more informative set of a genes will result in a similarity map that more accurately represents the true cell states. To compare the quality of different similarity maps, we made use of the surface protein abundance data which accompanies this single-cell mRNA-seq dataset as an independent indicator of cell state. We reason that that since protein abundance and mRNA expression are both derived from the cell state, an increase in the autocorrelation of protein abundance (when comparing two similarity maps generated from mRNA expresssion alone) indicates an increase in the quality of the similarity map.

The results of this evaluation (Figure 1B) show that, as we expected, employing feature selection procedures tended to result in increased protein autocorrelation when compared with a less-restrictive threshold based gene selection criteria. Furthermore, the procedure utilized by Hotspot showed greater increases than the other methods compared, though differences between the HVG, PCA, and Hotspot feature selection procedures were not statistically significant (ranksums test, p < .05).

After selection of the top 500 significant genes by Hotspot, we ran the module identification procedure on these genes (Figure 1C). Module 1 (includes CCR7 and SELL) and Module 3 (includes IL7R and S100A4) appear to distinguish the naive and activated T cell subsets while other modules illustrate transcriptional differences within the activated group. This includes Module 0 which highlights a subset of these cells expressing Th1 associated genes (including IFNG nd TBX21) and Module 2 which highlights another subset expressing higher levels of genes associated with cytotoxicity (PRF1, GZMB). Module 6, on the other hand, gathers genes associated with regulatory T cell activity (FOXP3, IL2RA and inhibitory receptors CTLA4, and TIGIT). In this manner, Hotspot is able to isolate distinct patterns of transcription even when they exhibit a nested, hierarchical structure.

### Combining Different Data modalities to identify Spatially-Dependent Gene Components

In the growing field of spatial transcriptomics ([29, 30, 11, 12]) new experimental methods have been developed which assay transcriptional profiles of single cells while also retaining information on their spatial origin. By using the these spatial positions to define the cell-cell similarity metric, Hotspot can be used to identify genes which drive spatial features such as spatially-dependent patterns of activation or non-random distributions of cell types.

To demonstrate this, we applied Hotspot to a sample consisting of 32,000 spatially-indexed transcriptional libraries from the mouse cerebellum [12]. As these libraries are sequenced at a low depth (median UMIs/barcode is 45), we ran Hotspot in ‘bernoulli’ mode where only the binary detection of a gene at each position is modeled (Methods). Based on their spatial distributions, 560 genes were identified with significant (non-zero) spatial autocorrelation (FDR < .05). These genes are enriched in marker genes for cerebellar cell types (Figure 2b) and are distinct from genes selected on the basis of high variability (Supplementary Figure 2A).

**Figure 2:**
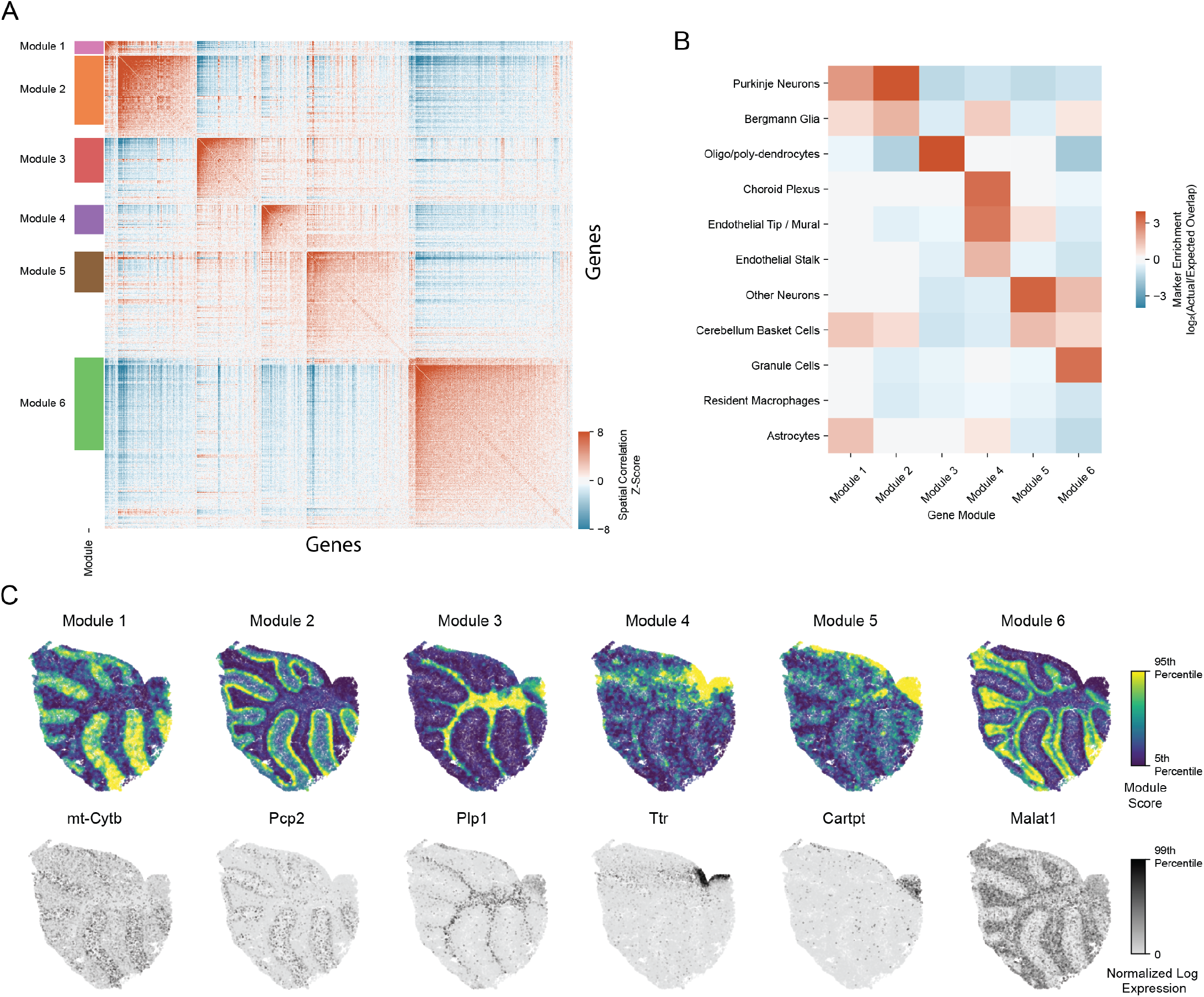
Spatial Gene Signatures Hotspot is used to identify spatially-relevant genes within a spatial single cell expression sample from mouse cerebellum. A) Genes with significant spatial autocorrelation (845 genes, FDR < .05) are grouped into 6 gene modules on the basis of pair-wise spatial correlations. B) Spatial modules are associated with specific cerebellar cell types as shown by enrichment in cell-type specific marker genes. C) Spatial gene modules are visualized with their summary, per barcode, module scores (top row). Beneath each plot of the module score, the expression of the gene with the highest spatial autocorrelation is visualized for comparison.

We then used Hotspot to group the genes into modules on the basis of pair-wise local correlation. This results in 6 transcriptional modules which correspond to known cerebellar cell types (Figure 2) and are reproducible between different cerebellar samples (Supplementary Figure 3). These modules reflect the primary structure of the cerebellum, with Module 1 capturing the Purkinje layer of neurons adjacent to the Granular layer (Module 6) followed by the oligodendrocytes of the white matter (Module 3). By computing summary module scores, the structure of the cerebellum becomes clearly visible (Figure 2C).

We compared our findings with an existing method for identifying expression patterns in spatial transcriptomics - SpatialDE [18]. A key difference between our method and SpatialDE is that the latter uses a Gaussian process model which requires the inversion of a large matrix as part of its optimization procedure. As a result, Hotspot is able to run much more quickly as the number of cells/positions increases (see comparison in Supplementary Figure 2B). To compare the results produced by each algorithm, we evaluated each for its ability to detect known positives and for its reproducibility between the four cerebellum samples produced by [12]. In using a set of marker genes for cell types in the mouse cerebellum (derived from [31]), we show that both methods achieve similar AUPR scores (Supplementary Figure 2C). To evaluate reproducibility, we used the IDR metric [32] to compare results between pairs of cerebellum samples and show that while both methods typically highlight several hundred genes at an IDR level of 0.1, the results produced by Hotspot are more consistent between pairs at this, and other, IDR thresholds (Supplementary Figure 2D).

In a similar manner, spatialDE is also able to identify modules of genes with similar spatial distributions of transcription. In comparing the gene modules output by Hotspot, we show that both methods are able to uncover similar patterns of variation (Supplementary Figure 4), with Hotspot requiring a much smaller runtime (93 seconds vs. 6.2 days for 10,000 cells).

### Identifying developmental expression in mouse embryogenesis

To demonstrate Hotspot on other measures of relatedness, we turned to a dataset of lineage-traced embryogenesis [14]. In this system, mouse embryos were engineered with a CRISPR Cas9 lineage-tracing system in which irreversible mutations are generated randomly throughout development at specified cut cites. The embryos then underwent single-cell sequencing at day 8.5 of development. Using the induced mutations, a cell’s developmental relationship to other cells can be assessed, and here we use this relationship to compute the cell-cell similarity metric when running Hotspot (see Methods for details). In this way, Hotspot can be used to extract genes whose expression is similar among similarly related cells and derive unsupervised modules associated with developmental changes.

We ran this procedure on 1756 cells from [14] and identified 2554 developmentally associated genes (FDR < .05). We further grouped these genes into 5 modules on the basis of expression changes with related cells (Figure 3A). To identify annotations for these modules, we made use of the annotated developmental cluster profiles from a separate dataset in [14] (Figure 3C. From this comparison, it is clear that module 1 (which includes markers Cubn, Amot, Amn, and Slc39a8) describes expression associated with the visceral and definitive endoderm, module 3 (with Hbb-bh1, Gata1, Klf1, and Nfe2) is associated with primitive blood, module 4 (with Srgn) is associated with the development of the parietal endoderm, and module 5 (with Plac1 and Ascl2) is associated with the differentiation of trophoblasts.

**Figure 3:**
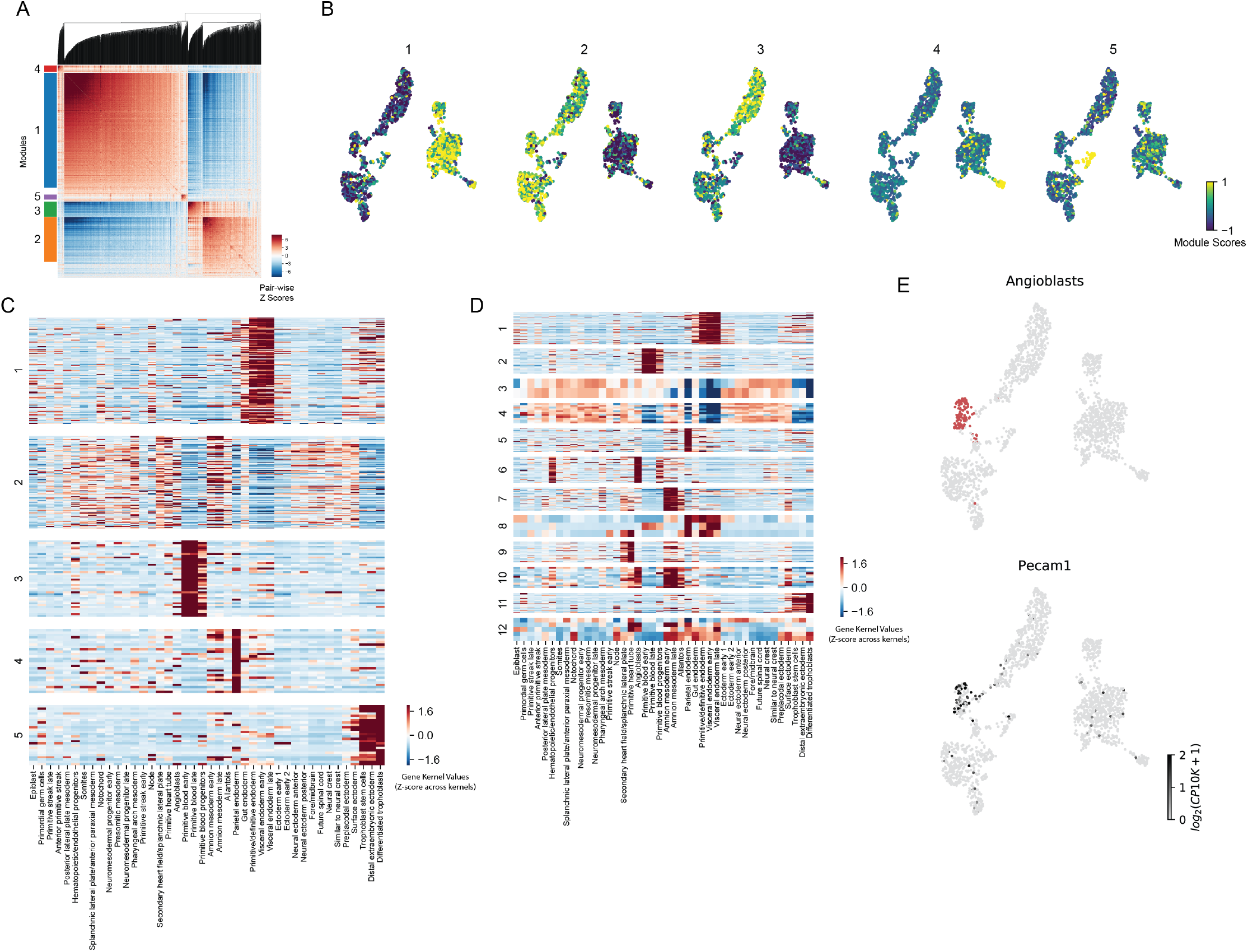
Lineage Gene Signatures. A) 2554 genes with significant (FDR < .05) lineage autocorrelation are grouped into 5 gene modules on the basis of pair-wise lineage correlation. B) Module summary scores are plotted against a UMAP of the expression data. C) A set of 42 manually annotated kernels from [14] representing cell types in mouse embryogenesis is used to identify putative labels for lineage-derived gene modules. For each module, the kernel values of the module genes are visualized. D) Similar to (C), only modules derived from gene expression (not lineage) are shown instead. E) Annotating cells which are most similar to the Angioblast kernel (top) along with an Angioblast marker gene, Pecam1 (bottom). Panels C-E show Hotspot is able to distinguish between gene modules with transcriptional support (such as that of Angioblasts) and those with lineage support.

Notably, an expression signature associated with the emergence of Angioblasts is identifiable when analyzing the expression data alone (Figures 3D, E) with marker genes such as Pecam1 having high autocorrelation in expression space. This same signature, however, is not detectable based on lineage similarity, implying that either emerging Angioblasts are less related than cells of other cell types at this developmental stage, or the number of cells or frequency of lineage-tracing mutations were insufficient to capture this effect. In this way, Hotspot can distinguish between genes and gene modules that have significant variation due to gene expression correlations, and those that arise from the inferred developmental tree.

## Discussion

Here we have described a set of techniques for feature selection and module identification in multi-modal single cell expression datasets. We demonstrate the application of these techniques using several previously published datasets. First, we show our approach may be used to identify spatially-varying gene expression and derive spatial gene modules using a single cell mouse cerebellar dataset [12]. Notably, we compare against a leading method designed specifically for this purpose, SpatialDE [18], and demonstrate comparable performance despite orders of magnitude of improvement in runtime. Second, in a dataset consisting of a combination of gene expression and lineage-tracing data [14], we demonstrate our approach for the purpose of identifying genes which exhibit lineage-dependent expression patterns. Finally, we demonstrate the utility of our approach when used with gene expression measurements alone. We use a combination of public data and simulated data to show that our approach is able to better identify relevant features than leading approaches (such as the commonly used highly-variable genes procedure) and that our metric of local correlation is able to better detect gene-gene correlations when compared with standard Pearson’s correlation.

The core of our method is the definition of two key test statistics. We defined a statistic for local autocorrelation within a KNN similarity graph that takes inspiration from the Geary’s C [23] and the Laplacian Score [24] which have been proposed for similar purposes. Notably our statistic differs from these by using a parametric null model for a gene’s expression to avoid the need for a permutation null and increase computational efficiency. Furthermore, we propose a novel pairwise local correlation statistic as an extension to this approach so that features (genes) may be grouped into modules on the basis of local expression similarity within the graph.

In addition to the examples shown here, the flexibility of our approach permits its application in other ways. Here we have varied the data used in the cell-cell metric (spatial, lineage, or transcriptional) and used Hotspot to identify genes with associated expression. However, the inverse is also possible in which the cell-cell metric is computed from gene expression and Hotspot is used to identify additional features that associate in expression space: for example, identifying specific lineage-tracing mutations, open chromatin regions (with combined single-cell expression and ATAC-seq [13]), or CRISPR perturbations associated with expression state changes. We have made the software behind the Hotspot procedure available at http://www.github.com/yoseflab/hotspot so that as single-cell technology evolves and additional modalities are incorporated, this framework can continue to be used extract signals across different classes of biological measurements.

## Methods

### A test statistic for feature selection

In analyzing single-cell RNA-seq with Hotspot, the first step consists of selecting genes which are informative, given a cell-cell similarity metric. Intuitively this can be thought of as identifying genes whose expression, if plotted onto a visualization of the cell manifold (such as those produced by tSNE or UMAP), produce non-random visual patterns and are therefore likely of interest to the analyst. In our previous work [33], we made use of the Geary’s C [23] as a test statistic to select gene signatures whose aggregate signature scores exhibited this property. One drawback with this statistic, however, is the lack of a well-defined null distribution, necessitating the use of permutations for significance testing. To eliminate the computational burden this produces when operating at the level of individual genes, we modify the Geary’s C and use…

The Geary’s C is defined as:

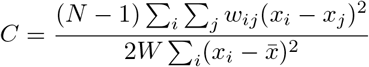

When applying this to scRNA-seq data, *N* represents the number of cells, *x_i_* represents the expression of the gene under test in cell *i*, and *w_ij_* represents the ‘weight’ between cells *i* and *j* (with *W* defined as the sum of all such weights over all cells). Weights are strictly non-negative and defined such that larger values are assigned to cells with higher similarity, with values decaying to 0 for highly dissimilar cells. In this way, lower values of *C* occur when similar cells tend to also have similar expression values for the gene being tested. It should be noted that a similar approach, termed the *Laplacian Score* [24], has been proposed for the use of feature selection for machine learning.

Here we modify this test statistic by removing terms which don’t specifically capture interactions between cells.

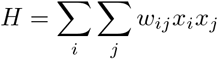

The removed terms in the numerator and denominator control for global distributional properties of the gene expression vector, *x*. Instead, we control for this by evaluating expectations of H and computing a Z-score as 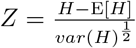.

Additionally, we assign weights using a K-nearest-neighbors graph, such that *w_ij_* is only positive if cells *i* and *j* are neighbors and there are no self-edges. As a result, the double summation can be re-expressed as a sum over edges, *E* in the resulting sparse graph:

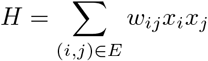

To evaluate expectations of H, a null model is needed. For our null model, we assume that expression values are drawn independently from some underlying distribution for which we can compute E[*x_i_*] and 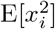 for each cell. Notably, values for E[*x_i_*] and 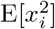 in the null model are estimated on a per-cell basis so that the effect of varying sequencing depths can be incorporated. Then expectations of H can be expressed as:

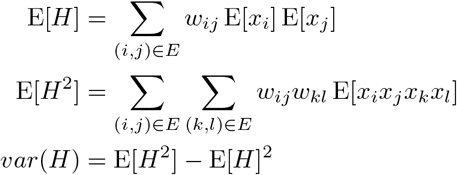

In computing E[*H*^2^], the lack of self edges implies that *i ≠ j* and *k ≠ l*, but the inner expectation may not be split as edge (*i, j*) can share nodes with (*k, l*). Additionally, computing E[*H*^2^] in this manner is difficult as a double summation over edges involves many terms (*O*(*N*^2^*K*^2^) for *N* cells and *K* neighbors). However, an alternate summation may be used that can be evaluated in *O*(*N K*) steps which first computes E[*H*^2^] assuming no edges share nodes and then corrects the (much fewer) terms for which this is not true:

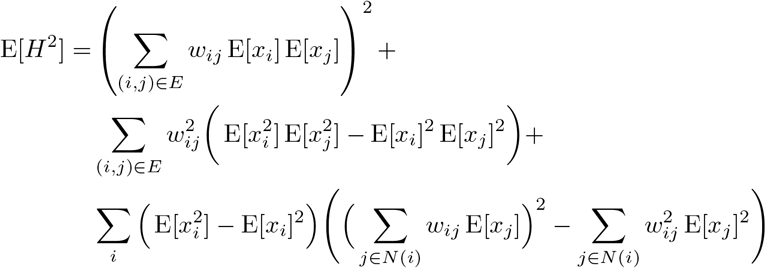

Finally, we can simplify further by standardizing each cell’s null model prior to computing *H*. Here we define:

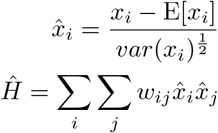

Computing the null model for 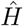 is then simplified as 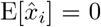 and 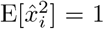 for all *i*. The resulting expression is then

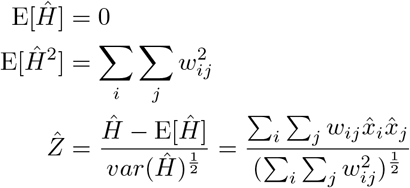

To compute p-values the resulting 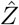 values are compared to the normal distribution. When testing multiple genes, the Benjamini-Hochberg procedure [34] is used to estimate the FDR.

### Null models for gene expression

In computing the autocorrelation, a model is needed for each gene and cell describing the expected distribution of expression values under the null hypothesis that each value is drawn independently. One utility of the approach described above is that only the first and second moments of this distribution are needed, allowing for flexibility in the choice of null model model. In this work, two different models were used and are described here. Importantly, both explicitly account for the per-barcode library size and adjust expected expression levels accordingly. This is necessary so that genes are not incorrectly flagged as significant due to autocorrelation in the library size.

#### Negative Binomial

The negative binomial distribution is a common choice to model the counts arising in single-cell RNA-seq experiments with UMIs [35]. Here we utilize the NBDisp model proposed in [27] in which the mean of the distribution for every gene is assumed to vary linearly with the library size for each cell. The model is defined as:

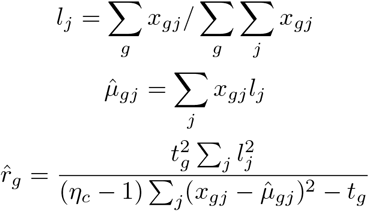

for gene *g* and cell *j*. *t_g_* represents the total number of UMI counts for gene *g, η_c_* is the total number of cells, and 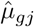 and 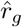 represent the negative binomial mean and dispersion. Moments of expression are then estimated as:

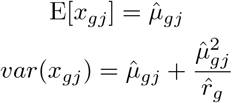

#### Bernoulli

As an alternative to the negative binomial model, we define a “Bernoulli” model where only the *detection* of a gene (defined as a UMI count greater than 0) is estimated. This model may be a better choice for extremely sparse data where detecting more than one count per gene in a cell is rare (such as the Slide-Seq [12] analyzed in this article.) The model for gene *g* is formulated as:

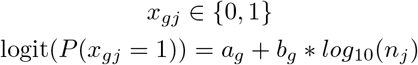

where *n_j_* is the total number of UMI in cell *j*. Model parameters *a_g_* and *b_g_* are fit on a per-gene basis. First cells are aggregated into 30 bins based on UMI, with bin center’s 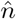 and the average probability of detection per bin is computed as 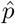. Then linear regression is used to estimate model coefficients as 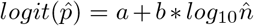. We considered the use of logistic regression directly on the sample values, but due to performance issues decided to utilize this binned approximation.

### Deriving gene modules

To compute gene modules, Hotspot uses a three-step procedure:

1. Find informative genes with high local autocorrelation
2. Evaluate pair-wise local correlations between genes
3. Cluster the resulting gene-gene affinity matrix

For step (1), the feature selection procedure described above is applied and a cutoff is used to select the most highly-informative genes. In the step (2), we modify our procedure for feature selection to evaluate correlations between genes in a manner that also leverages the global cell-cell similarity map. We denote this ‘local correlation’. In this way, gene pairs which tend to be sparsely expressed in the same regions of the similarity map can be detected as correlated even if they are infrequently detected in the same cell. Finally, step (3) involves in clustering the genes by genes local correlation matrix computed in step (2). Here we found good performance in running a modified hierarchical clustering procedure (details follow).

#### Evaluating Pair-Wise Local Correlation

We define the following to evaluate local correlations between genes *x* and *y*:

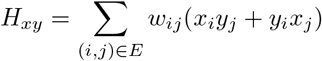

Notably, while local autocorrelation tends to only exhibit positive values, local correlations will often produce negative values. This makes sense intuitively as genes expressed in different regions of the similarity map will overlap *less* than expected if there values were drawn at random.

For the purpose of clustering, we transform this local correlation into a Z-score by comparing it to its expected first and second moments under a null model. We initially considered a null model that assumes expression values for genes *x* and *y* are all independent. However, this formulation tends to significantly underestimate the variance of the test statistic if at least one gene has high *autocorrelation* - which is a guarantee since we pre-select these genes. Instead, we compare against a null model where one gene’s values are fixed and the other are assumed to be independent. In other words “given the observed of gene *x*, how extreme is *H_xy_* compared with independent values of *y*”. Since the test statistic is symmetric with respect to the choice of *x* and *y*, we compute Z-scores using both *P*(*H_xy_|x*) and *P*(*H_xy_|y*) and conservatively retain the least-significant (closest to zero) result.

The moments of *H_xy_* under *P*(*H_xy_|x*) are computed as:

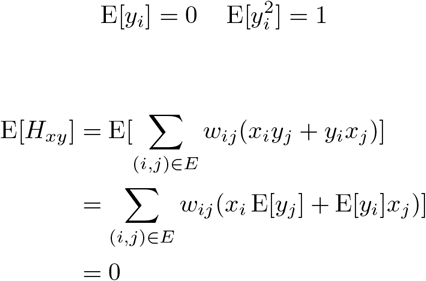

Values of *y* have been standardized with respect to one of the previously described gene models. Values of *x* are held fixed (i.e., they are not random variables).

Computing 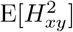 is more involved. First we note that the value can be expressed as a sum of edge pairs:

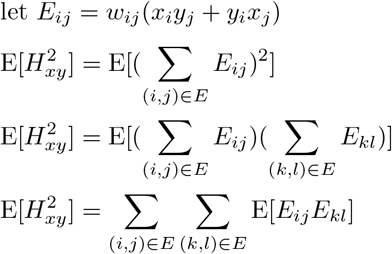

To compute this efficiently, we make note that when evaluating expectations of pairs of edges, E[*E_ij_E_kl_*], there are three possible situations:

1. edge pair shares no noes (*i, j, k, l* all distinct):

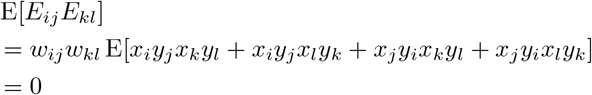
2. edge pair shares one node (for example, *i = k*):

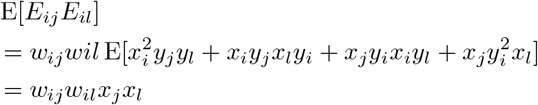
3. edge pair shares two nodes (for example, *i = k* and *j = l*):

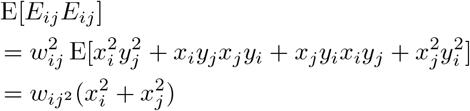

Since only products of edges that share neighbors need to be considered, we can compute the expectation in *O*(*E*) time as:

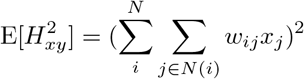

where *N* is the number of nodes and *N*(*i*) are nodes which share an edge with node *i*.

#### Clustering the Gene-Gene Affinity Matrix

Once the gene by gene matrix of Z-scores has been computed, we apply a bottom-up clustering procedure with two parameters: MIN_CLUSTER_GENES,and FDR_THRESHOLD. Iteratively, the two genes/modules with the highest pair-wise Z-score are merged, using the UPGMA procedure to derive updated Z-scores between the resulting module and the remaining modules and/or genes. If a module accumulates more than MIN_CLUSTER_GENES, then it is assigned a label. To preserve hierarchical structure between modules, if two labeled modules are merged, a new label is not assigned and genes merged into the resulting composite module remain unlabeled. The FDR_THRESHOLD parameter is used to set a minimum significant Z-score by applying the Benjamini-Hochberg procedure [34] to the associated p-values of all Z-scores and selecting the minimal such Z-score which is below the FDR threshold. If at any point in the above merging procedure the maximal Z-score falls below this threshold, the procedure halts as further gene assignments fall below the significance threshold and are therefore ambiguous.

#### Computing Per-Cell Module Scores

To visualize gene modules, it is useful to evaluate per-cell module scores which can then be plotted onto UMAPs, spatial plots, dendrograms, etc. When computing module scores, Hotspot uses the following procedure: first counts are centered using the selected null model (e.g., ‘bernoulli’ or ‘Negative Binomial’). Then the resulting values are smoothed using the KNN graph - the smoothed expression for each cell is computed as the weighted average of its neighbors in the graph. These smoothed values are then modeled with PCA using a single component, and the cell-loadings are reported as the model scores (potentially with a sign flip so that gene coefficients are positive).

### Analysis of Mouse Cerebellum samples

Barcode expression and position information was downloaded for four mouse cerebellum samples (Puck_180819_9, Puck_180819_10, Puck_180819_11, and Puck_180819_12) from https://portals.broadinstitute.org/single_cell/study/slide-seq-study as directed in [12]. For all analyses involving a single sample, Puck_180819_12 was used.

When running Hotspot, genes were initially prefiltered to remove those detected in less than 50 cells. For feature selection, a neighborhood size of 300 was used and the ‘bernoulli’ model was used to model gene detection probabilities. For pair-wise correlation and module identification, genes were selected with an FDR < .05 and the neighborhood size was reduced to 30 to identify spatial modules at a finer resolution.

When running spatialDE, genes were again prefiltered to remove those detected in less than 50 cells. Initial runs of spatialDE tended to erratically select an unreasonably small length scale with poor results, and so we recomputed the length-scale values with a minimum of 50 prior to fitting the model. When computing modules with spatialDE, we selected a length scale of 350 (following the tools guidelines to select a length scale slightly larger than the average per-gene optimal length scale) and 10 components as an initial run with 5 components did not appear to capture all spatial patterns.

To evaluate the performance of feature selection with the cerebellum data, we sought to create a list of ‘true’ positives. We reasoned that the spatial patterns in the cerebellum would be largely influenced by changes in the composition of cell types and so marker genes for cerebellum cell types could be used a positive gene list. To create this list, we downloaded raw expression data from the DropViz [31] database and computed marker gene sets for each of the 11 annotated mouse cerebellum clusters. For each cluster, we evaluating a 1 vs. all ranksums test for differential expression on every gene and retained the top 100 genes above a significance threshold (FDR < 0.1). The eleven lists were then merged to create a true positive set for the precision-recall curves. These same marker gene lists were also used when characterizing Hotspot modules.

In addition to this marker-gene based approach, we evaluated performance based on reproducibility between pairs of mouse cerebellum samples. For the four samples, Hotspot and spatialDe were both run in the same manner as described above and for each of the six sample pairs, the reproducibility of the gene ordering was evaluating using the Irreprodicible Discovery Rate (IDR) metric [32].

### Analysis of Mouse Embryogenesis Lineage-Tracing Data

Expression data from Chan et al. [14] was downloaded from NCBI GEO (accession GSE117542). For this analysis, the sample corresponding to Embryo 3 was used. The developmental lineage was inferred by running the Cassiopeia-Greedy algorithm [36] with priors determined from the frequency of indels across all observed embryos (as done in the original study [14]). Hotspot was run using two different metrics to compare outputs: the ‘tree’ metric in which the KNN graph was formed from the 30 nearest neighbors according to the inferred lineage and the ‘transcription’ metric in which PCA was run (on *log*_2_(*x* + 1)-transformed counts-per-10,000 expression values) and the KNN graph was formed from the 30 nearest neighbors in the reduced (top 20 component) transformed space. For both cases, genes were pre-filtered to remove those expressed in less than 10 cells, and the negative binomial model was used (with adjustment for library size). To evaluate local correlations between genes, for the tree-metric all genes with autocorrelation passing the 0.05 FDR threshold were selected. For the transcription metric, a large number of genes has significant autocorrelation and so the top 2000 (based on Z-score) were selected for local correlations. When extracting modules, a local correlation FDR threshold of 0.05 was used and `MIN_CLUSTER_GENES` settings of 30 and 50 were used (for tree-metric and transcription-metric respectively).

In evaluating the modules computed from Hotspot, the expression data from [14] was used. Specifically the Cell State Kernels were downloaded from NCBI GEO (accession GSE122187) and for each module returned by Hotspot, the kernel values were plotted (after standardizing across cell types) for the intersection of the 712 kernel genes and the genes in each module.

UMAP projections were generated by first running PCA (as described above) and then running UMAP [22] on the transformed space (top 20 components) with `n_neighbors=30`.

### Analysis of CD4 T Cell Transcriptomes

For the CD4 T cells, we made use of the public “5k_pbmc_protein_v3” available from 10x Genomics. Digital gene expression was downloaded from 10x and the full set of PBMCs was filtered based on surface protein abundance to extract CD4 T cells. First, the protein abundance measurements were log-transformed (*log*(*x* + 1)) and then each protein was mean-centered individually. Cells were retained based on manual thresholding of bimodal markers (specifically CD4 > 3, CD3 > 2.5, CD14 < 2). Additionally, we discarded cells with low recovered mRNA UMI counts (UMI < 3100) or with a high proportion of counts from mitochondrial genes (>16%). This resulted in 1486 CD4 T cells.

When running the Highly Variable Genes procedure, Seurat’s *FindVariableGenes* method was used. First counts were log-normalized with a scale factor of 10,000. Then *FindVariableGenes* was run with a x.low.cutoff = .01, x.high.cutoff = 3.04 (corresponds to a high-end cut-off of 20 counts/10,000), and y.cutoff = .5. Resulting genes were then ordered in descending order by the reported ‘gene.dispersion.scaled’.

For the NBDisp procedure, the NBumiFitModel function as implemented in the M3Drop [27] package was used with default settings on raw gene counts. Additionally, for consistency with the HVG procdure, a high-end cutoff of 20 counts/10,000 was used. Output genes were ordered by the reported ‘q.value’ in ascending order.

For the PCA procedure, the feature proposed by [27] was used. Specifically, principal components analysis was run with 5 components on the log-transformed, scaled counts/10,000. A genes score was reported as the sum of the absolute value of its PCA coefficients in the top 5 components. Genes were then ordered in descending order based on this score.

To compute gene relevance (GR) scores, we downloaded the c7 immune signature set from MSigDB [28]. Gene sets were filtered to retain sets describing comparisons between CD4 T cells, specifically sets with one of the following terms (Th1, Th2, Tfh, Treg, Tconv, T, Th17, Tcell, NKTcell, CD4, Thymocyte) and none of these terms (DC, PDC, Macrophage, BMDM, Monocyte, Neutrophil, Mast, B, BCell, NKCell, NK) resulting in 706 CD4-relevant gene sets. For every gene, its GR score was computed as the number of CD4-relevant gene sets in which it appears. When evaluating a set of genes (selected by the Hotspot, HVG, NBDisp, or PCA procedure), the resulting metric was reported as the average GR score for the set.

To evaluate a set of genes using protein autocorrelation, the raw counts were subset to include only the genes under evaluation and then scVI [16] was run with 10 latent components. Hotspot was then used to evaluate the autocorrelation of the surface protein measurements. This is predicated on the assumption that a more informative set of genes leads to a latent space model that more accurately reflects true cell state, and that both mRNA expression and protein surface abundance are derived from this cell state. Therefore, we would expect a more informative set of genes to result in increased autocorrelation values for surface proteins as well. The resulting protein Z-scores for each of the four methods were then compared with the Z-scores from a fifth run where genes were selected with a simple thresholding procedure (all genes expressed in at least 10 cells).

Other implementation details are as follows: Within Hotspot, protein abundance was modeled using a depth-adjusted normal model. First protein counts were log-transformed, and then the log of the total protein counts (per cell) were linearly regressed out of the transformed protein counts as we observed significant correlation between individual protein counts and the total counts in other proteins, per cell. This normalized protein abundance matrix was then standardized on a per-protein basis. In selecting genes with Hotspot, PCA was first used with 20 components to create the latent space for cell-cell similarities. For all procedures, the top 1000 reported genes were used. To avoid evaluating on surface proteins which were not expressed in CD4 T cells (and whose detection therefore represents background noise), we fit a 2-component gaussian mixture model on the log-transformed surface abundance measurement on all PBMCs for each protein and retained proteins where the sparation between peaks (in log-space) was at least 2 and at least 1% of CD4 T cells were assigned to the higher (active) component.

In identifying expression modules in the CD4 T cells, first SCVI was run on the top 1000 highly variable genes with 10 components. This latent space was then used as input to Hotspot and modules were created from top 500 genes based on autocorrelation. For module creation, an FDR_THRESHOLD of 0.05 was used along with a MIN_CLUSTER_GENES setting of 15.

### Simulated Transcriptional Profiles

Simulated data was used to evaluate the performance of Hotspot when run on transcriptional data alone. To generate simulated single-cell profiles, we used the SymSim [25] framework. For each of 10 simulation replicates, UMI count profiles were generated for 3000 cells consisting of 5000 genes. 5 gene components were generated by varying the ‘synthesis’ parameter with 100 genes participating in each component. The components were generated with the following structure:

- Component 1: Affects all cells
- Component 2: Affects 300 cells which had positive Component 1 values
- Component 3: Affects 30 cells which had positive Component 1 values
- Component 4: Affects all cells which had negative Component 1 values
- Component 5: Affects all cells

For full implementation details, see accompanying code.

When evaluating feature selection on this data, methods were run as described here. Seurat’s [26] ‘FindVariableGenes’ function was used as the Highly Variable Genes (HVG) method with *x*.low.cutoff = 0.1, *x*.high.cutoff = 100, and y.cutoff = 0.5. The NBDisp method from [27] was run with default settings. For the PCA procedure, as described in [27], we first ran PCA on the *log*(*x* + 1) transformed scaled counts per 10,000 values retaining the top 5 components. Genes were then ordered using the sum of the absolute value of their component weights in descending order. When running Hotspot, we used the cell-components of this same PCA procedure as the latent space to construct the KNN graph (300 neighbors), and the NBDisp model (from [27]) as the expression null model.

To evaluate the module assignment procedure, we sought to compare whether the use of Hotspot’s local correlation resulted in an improvement over Pearson’s correlation - the intuition being that taking into account a cell’s local neighborhood should increase the signal of an association between two genes with true, biological correlation. To decouple feature selection from this evaluation, we pre-selected the 500 non-random genes and computed both Hotspot’s pair-wise local correlation (Z-scores) and Pearson correlation coefficients between gene pairs. For evaluating Pearson correlation, the standardized values (under the library-size-adjusted negative binomial distribution) were used when computing correlation coefficients and associated p-values. Then, genes were grouped into modules using the hierarchical procedure implemented by Hotspot and an FDR cutoff of 0.05. The modules formed by either procedure clearly mapped clearly to the original 5 modules, and so we evaluated each by computing the overall accuracy (proportion of genes correctly assigned to their true module) over the 500 genes. Additionally, the accuracy was computed within bins of genes based on the gene mean expression and the gene-effect coefficient magnitude used by SymSim. As with the feature selection evaluations, this was repeated over 10 replicate simulations.

---

**Supplementary Figure 1:**
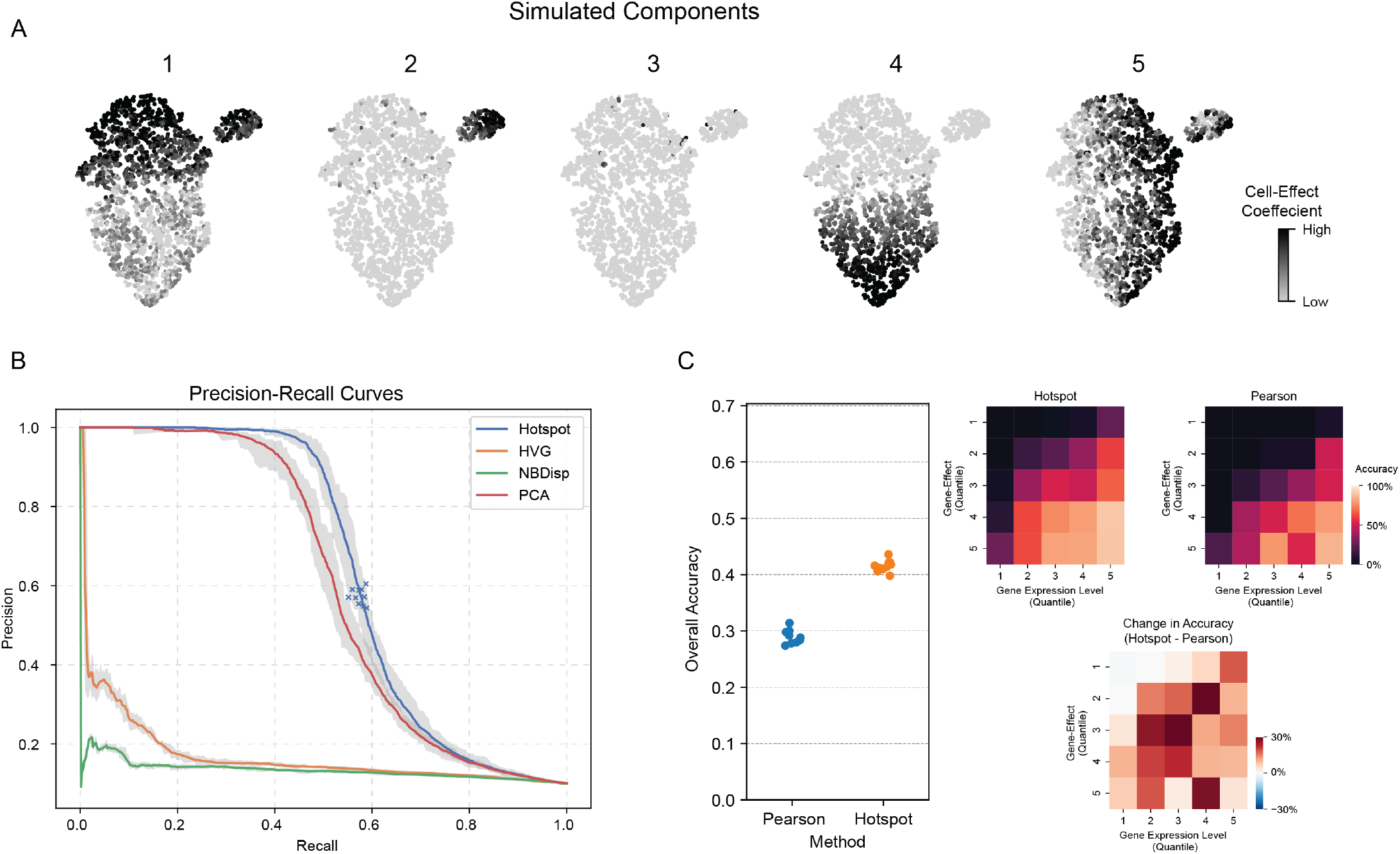
Evaluation with Simulated Transcriptional Profiles. A) SymSim was used to simulated 5 gene modules (each with 500 genes) affecting a varying number of cells (see Methods for full details). Visualized here are the cell-effect coefficients on a UMAP projection of the simulated dataset. B) Precision and Recall for four methods of feature selection - Hotspot, HVG (highly variable genes, as implemented in Seurat[26]), NBDisp (from [27]), and a PCA-based procedure (as described in [27]). Evaluations from 10 replicate simulations are shown with the trace denoting the mean and the shading denoting the minimum and maximum across replicates. Point-estimates are shown for Hotspot (crosses) when selecting an FDR threshold of 0.05. C) Comparison of the local correlation statistic developed here with that of Pearson correlation for the task of correctly assigning genes to the same module. Left panel shows overall accuracy while right shows accuracy per 5 gene-effect quantiles (quantile 1 represents genes with the weakest simulated effect strength, 5 the strongest). Bars represent the mean of 10 simulations and vertical lines denote bootstrapped 95% confidence intervals.

**Supplementary Figure 2:**
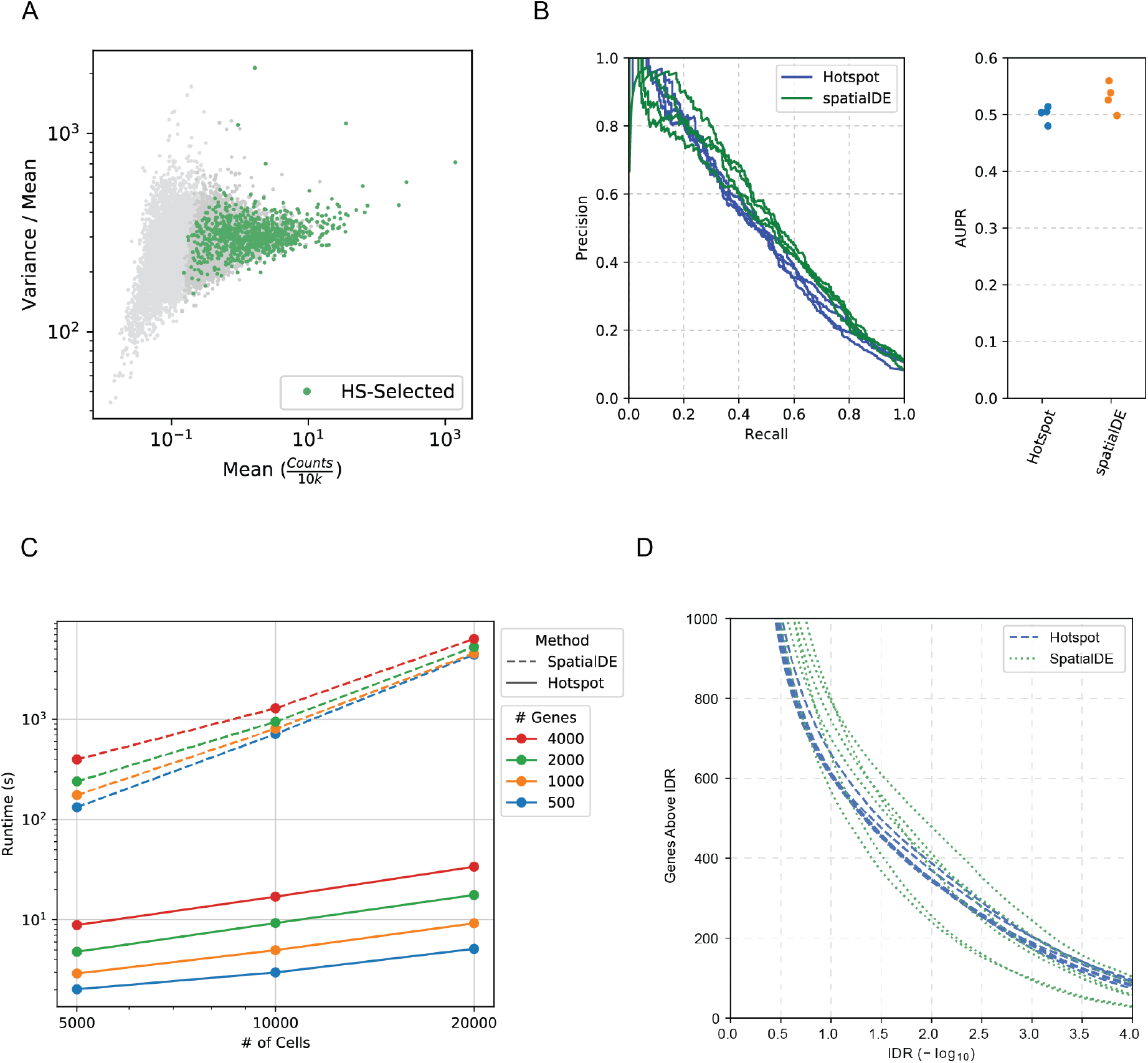
Supplementary Figures for Spatial Gene Signatures. A) Mean and Fano factor of expression measurement for all genes in the Puck 180819 12 mouse cerebellum sample. B) Comparison of SpatialDE and Hotspot in terms of their ability to identify marker genes for mouse cerebellar cell types. C) Runtime comparison between SpatialDE and Hotspot. D) Comparison of Hotspot and Spatial in terms of reproducibility between similar samples. The Irreproducible Discovery Rate (IDR) metric is used to compare genes selected between six pairs of mouse cerebellar samples.

**Supplementary Figure 3:**
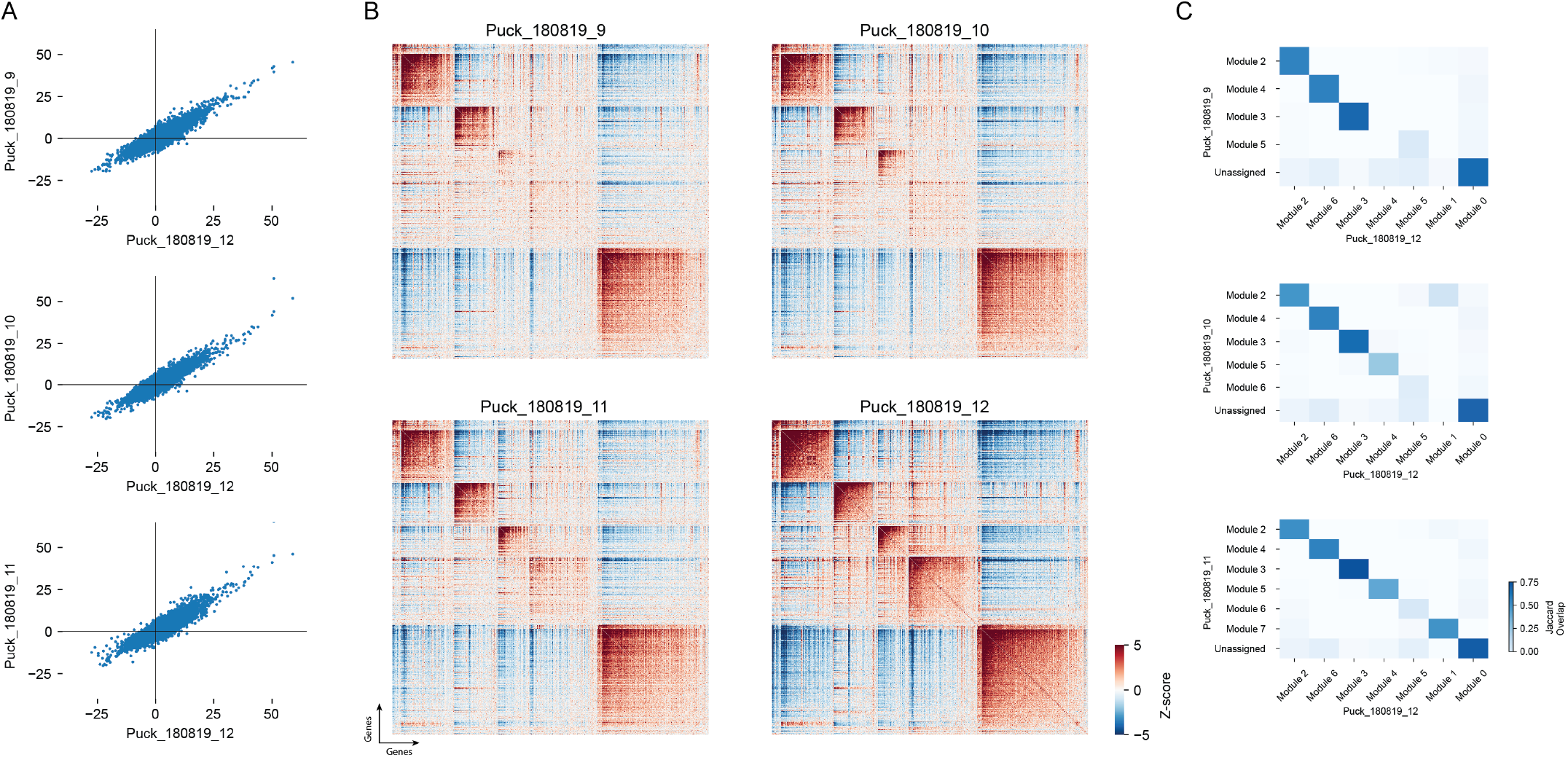
Comparison of spatial gene correlations between samples. A) For each pair of the 845 spatial genes (taken from the Puck_180819_12 sample), the spatial correlation Z-score is evaluated in all four samples and results are compared with that of sample Puck_180819_12. B) the same pair-wise Z-scores from (A) are visualized as a gene by gene correlation plot to compare the modules across the four samples. C) A comparison of the overlap between gene modules identified in the Puck_180819_12 sample and those identified in the other 3 samples.

**Supplementary Figure 4:**
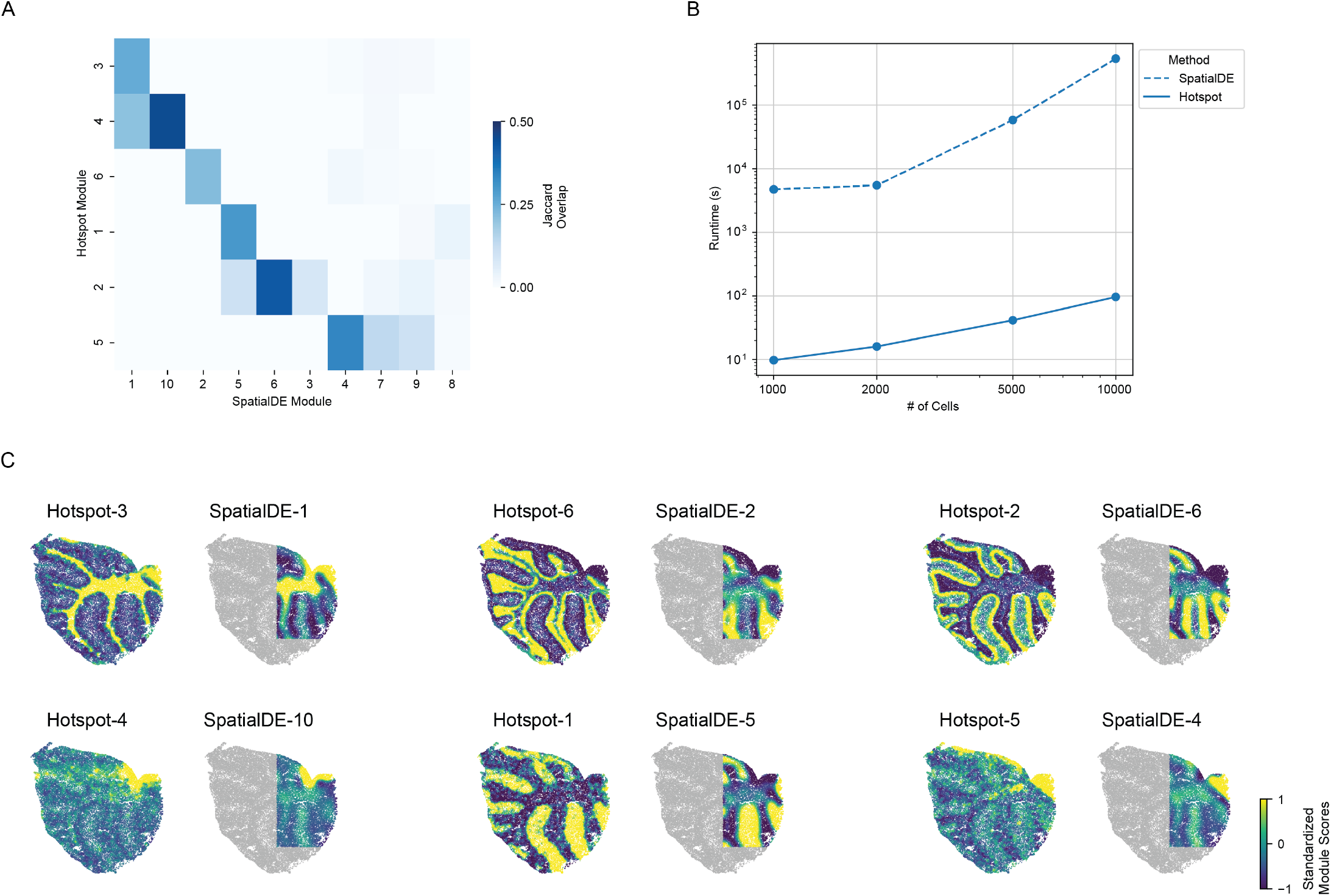
Comparison of modules identification procedure in Hotspot and SpatialDE. A) Overlap coefficient between modules identified by Hotspot and modules identified by SpatialDE in the Puck_180819_12 sample. B) Runtime comparison for the two algorithms with varying number of cells. Each trial run on 500 genes using 16 threads, one core per thread. C) Hotspot/SpatialDE module pairs (from A) are visualized using the spatial coordinates of the sample. Both procedures are able to identify similar patterns with Hotspot running in orders of magnitude less time.

**Supplementary Figure 5:**
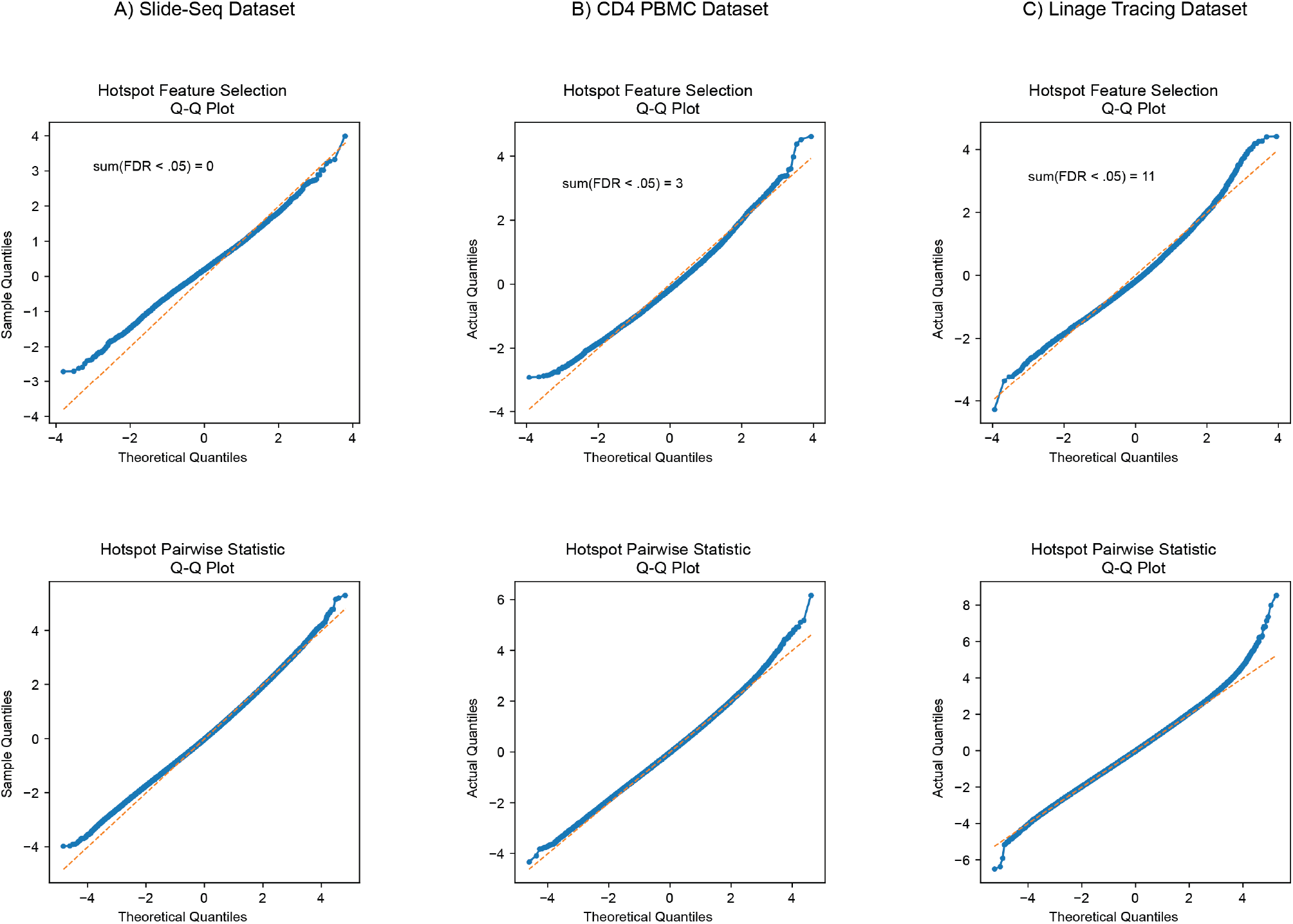
Q-Q plots for test statistics. For each dataset analyzed in the study, the theoretical quantiles of the test statistic (for local autocorrelation, top row, and for local pair-wise correlation, bottom row) are compared against those computed on shuffled data. A) Expression/positions from Puck_180819_12 of the Slide-Seq [12] data. B) CD4 Expression data from 10x Genomics. C) Expression and lineage data from [14], Embryo3. When computing shuffled Z-scores for pairs of genes, only the second gene in the pair is shuffled.

